# Pelagiphages in the *Podoviridiae* family integrate into host genomes

**DOI:** 10.1101/410191

**Authors:** Yanlin Zhao, Fang Qin, Rui Zhang, Stephen J. Giovannoni, Zefeng Zhang, Jing Sun, Sen Du, Christopher Rensing

## Abstract

The *Pelagibacterales* order (SAR11) in Alphaproteobacteria dominates marine surface bacterioplankton communities, where it plays a key role in carbon and nutrient cycling. SAR11 phages, known as pelagiphages, are among the most abundant phages in the ocean. Four pelagiphages that infect *Pelagibacter* HTCC1062 have been reported. Here we report 11 new pelagiphages in the *Podoviridae* family. Comparative genomic analysis revealed that they are all closely related to previously reported pelagiphages HTVC011P and HTVC019P, in the *HTVC019Pvirus* genus. *HTVC019Pvirus* pelagiphages share a core genome of 15 genes, with a pan-genome of 234 genes. Phylogenomic analysis clustered these pelagiphages into three subgroups. Integrases were identified in all but one pelagiphage genomes. Evidence of site-specific integration was obtained by high-throughput sequencing and sequencing PCR amplicons containing predicted integration sites, demonstrating the capacity of these pelagiphages to propagate by both lytic and lysogenic infection. HTVC019P, HTVC021P, HTVC022P, HTVC201P and HTVC121P integrate into tRNA-Cys genes. HTVC011P, HTVC025P, HTVC105P, HTVC109P, HTVC119P and HTVC200P target tRNA-Leu genes, while HTVC120P integrates into the tRNA-Arg. Evidence of pelagiphage integration was also retrieved from Global Ocean Survey (GOS) database, suggesting the occurrence of pelagiphage integration *in situ*. The capacity of *HTVC019Pvirus* pelagiphages to integrate into host genomes suggests they could impact SAR11 populations by a variety of mechanisms, including mortality, genetic transduction, and prophage-induced viral immunity. *HTVC019Pvirus* pelagiphages are a rare example of a lysogenic phage that can be implicated in ecological processes on broad scales, and thus have potential to become a useful model for investigating strategies of host infection and phage-dependent horizontal gene transfer.

**IMPORTANCE:** Pelagiphages are ecologically important because of their extraordinarily high census numbers, which makes them potentially significant agents in the viral shunt, a concept that links viral predation to the recycling of dissolved organic matter released from lysing plankton cells. Lysogenic Pelagiphages, such as the *HTVC019Pvirus* pelagiphages we investigate here, are also important because of their potential to contribute to the hypothesized processes such as the “Piggy-Back-the-Winner” and “King-of-the-Mountain”. The former explains nonlinearities in virus to host ratios by postulating increased lysogenization of successful host cells, while the latter postulates host-density dependent propagation of defensive alleles. Here we report multiple Pelagiphage isolates, and provided detailed evidence of their integration into SAR11 genomes. The development of this ecologically significant experimental system for studying phage-dependent processes is progress towards the validation of broad hypotheses about phage ecology with specific examples based on knowledge of mechanisms.

## INTRODUCTION

Viruses are extremely abundant in marine systems, where they outnumber plankton cells by approximately 10-to-1 (1). Recent reports have shown the relationship between virus-like particles and plankton cells follows a power law function, declining at high cell abundances (2, 3). Most marine viruses are bacteriophages that infect bacterial hosts (1, 4-7). Bacteriophages affect the microbial community composition and influence microbial community diversity in ocean ecosystems by a variety of mechanisms (1, 8). For example, the influential “Kill-the-Winner” hypothesis describes the impacts of density-dependent lytic phage predation (8-10), and the recent “Piggyback-the-Winner” model proposes that lysogeny by temperate phages is more prevalent when host populations are being successful (2). Phages are also known to drive host evolution by mediating genetic exchange and exerting predation pressures that favor continuous evolution at the population level and divergence at loci that encode resistance (11-13). In the “King-of-the-Mountain” hypothesis, we proposed that host cell density-dependent horizontal gene exchange could alter the frequencies of alleles for phage defense (14).

The ubiquitously distributed marine SAR11 bacteria (order *Pelagibacterales*) are considered as the most abundant and successful organisms in the world (15). They contribute significantly to marine biomass and oxidize marine dissolved organic matter oxidization, thus they have a major impact on the global carbon cycle (16). Given their ubiquity and abundance, SAR11 bacteria are the ideal targets for phage attachment and predation. However, considering the importance of SAR11, the study of phages infecting SAR11 bacteria (pelagiphages) has lagged considerably behind the study of other marine phages. Only four cultured pelagiphage genomes have been sequenced (14). Metagenomic recruitment indicated these pelagiphages are prevalent, constituting an important component of marine viral communities (14). Of the four pelagiphages, HTVC019P and HTVC011P are short-tailed podoviruses belonging to the *Autographivirinae* subfamily of the which was previously termed as T7-like phages (17). They share a common evolutionary origin with other *Autographivirinae* genera.

HTVC019P and HTVC011P were found to contain the integrase gene (14). Integrases carry out site-specific recombination between the phage chromosomal attachment site (*attP*) and the bacterial chromosomal attachment site (*attB*) that are markers for prophages (18). Short-tailed *Autographivirinae* phages are typically host-specific and previously have been known to have lytic life-cycle strategies. However, in 2002, a prophage was identified in the *Pseudomonas putida* KT2440 genome that had sequence similarities to phage T7 (19). This prophage was shown to be active, which implied that some *Autographivirinae* phages might be able to develop the lysogenic life strategies. Thereafter, more *Autographivirinae* prophage structures were identified in some bacterial genomes. In addition, integrase genes were identified in nine of twenty sequenced marine *Autographivirinae* cyanophage genomes (20-23), but no evidence shows these cyanophages have the ability to integrate their genomes into the host chromosomes. Cyanophage P-SSP7 carries an integrase, and 42 bp exact match to partial host tRNA-Leu, which is a putative phage integration site (20). With the emergence of cultivation-independent metagenomic sequencing techniques, representatives of novel marine pelagiphage genomic contigs belonging to *Autographivirinae* subfamily and other phage subfamilies were identified from Mediterranean DCM (MedDCM) fosmid library (24). Integrases and exact matches to SAR11 tRNA genes were also identified in some pelagiphage contigs (24), implying the prevalence of integrase in pelagiphage genomes.

This study reports an extensive research effort made to isolate and study additional HTVC019P-related pelagiphage representatives. Comparative genomics provided insight into conserved genomic features and evolutionary relationships of these pelagiphages. We propose to place these phages in a new genus, *HTVC019Pvirus*, in the *Autographivivinae* subfamily. Most of the *HTVC019Pvirus* pelagiphages were demonstrated to integrate into the SAR11 chromosomes in culture. Their exact integration sites were mapped by using nexy-generation sequencing and confirmed by PCR amplification of phage integration sites.

## RESULTS AND DISCUSSION

### General features of *HTVC019Pvirus* pelagiphages

A total of 11 short-tailed pelagiphages were isolated from diverse marine environments using three closely related SAR11 strains that belong to SAR11-Ia group (98.9-99.4% 16s rDNA sequence identity). These pelagiphages were tested for their ability to infect three SAR11 hosts. All pelagiphages isolated from HTCC1062, two isolated from HTCC7211 and one isolated from FZCC0015 only infected their original hosts, while HTVC200P, HTVC121P, HTVC119P and HTVC109P infected both HTCC7211 and FZCC0015 (Table 1).HTCC7211 and FZCC0015 are more closely related (99.4% 16S rDNA sequence identity; ANI=80.38%). Generally, podoviruses possess narrow host ranges. The cross-infection results indicated that these pelagiphages likely have narrow host ranges, could only infect very closely related hosts.

**Table 1.** General features of pelagiphages analysed in this study.

| Phage | Original host | Host infectivity |  |  | Source water | Depth | Genome size<br>(bp) | G+C<br>% | NO. of<br>ORFs | Accession number | Reference |
| --- | --- | --- | --- | --- | --- | --- | --- | --- | --- | --- | --- |
|  |  | HTCC1062 | HTCC7211 | FZCC0015 |  |  |  |  |  |  |  |
| HTVC021P | HTCC1062 | + |  |  | South China Sea K2 | 5 m | 42,809 | 33.5 | 60 | MH579717 | This study |
| HTVC022P | HTCC1062 | + |  |  | South China Sea A9 | 20 m | 42,010 | 34.2 | 54 | MH598798 | This study |
| HTVC025P | HTCC1062 | + |  |  | Baltic Sea | surface | 37,251 | 32.5 | 49 | MH598799 | This study |
| HTVC031P | HTCC1062 | + |  |  | Osaka Bay, Japan | surface | 41,046 | 33.4 | 53 | MH598700 | This study |
| HTVC201P | FZCC0015 |  |  | + | Yantai coast, Bohai sea | surface | 41,415 | 33.1 | 52 | MH598802 | This study |
| HTVC200P | FZCC0015 |  | + | + | Osaka Bay, Japan | surface | 42,221 | 33.2 | 53 | MH598801 | This study |
| HTVC121P | HTCC7211 |  | + | + | South China Sea K3 | 150 m | 42,600 | 33.5 | 57 | MH598803 | This study |
| HTVC105P | HTCC7211 |  | + |  | Indian Ocean I109 | 75 m | 42,385 | 33.5 | 56 | MH598804 | This study |
| HTVC109P | HTCC7211 |  | + | + | BATS Hydrostation S<sup>a</sup> | 200 m | 41,323 | 35.5 | 54 | MH598805 | This study |
| HTVC119P | HTCC7211 |  | + | + | Yantai coast, Bohai sea | surface | 38,357 | 32.0 | 54 | MH598806 | This study |
| HTVC120P | HTCC7211 |  | + |  | Yantai coast, Bohai sea | surface | 42,622 | 33.3 | 53 | MH598807 | This study |
| HTVC019P | HTCC1062 | + |  |  | Oregon NH10 | 10 m | 42,101 | 34.0 | 59 | KC465901 | Zhao et al., 2013 |
| HTVC011P | HTCC1062 | + |  |  | Oregon NH10 | 10 m | 39,921 | 32.0 | 45 | KC465900 | Zhao et al., 2013 |

a. BATS: Bermuda Atlantic Time Series Station

The general features of the pelagiphages analyzed in this study are shown in Table1. The 11 newly isolated pelagiphages have genome sizes of between 37.2 kb and 42.8 kb, coding for 45-60 ORFs, similar to the genome size and number of ORFs of HTVC019P and HTVC011P. The G+C% of these 11 pelagiphages range from 32.0% to 35.5%, significantly lower than G+C% of other *Autographivirinae* phages, but similar to those of their hosts (29.03-29.68%) and previously reported pelagiphages (29.7-34.0%) (14). Approximately 80-90% of the ORFs share a significant similarity to previously identified gene products from diverse organisms. Approximately 40% of all identified ORFs can be assigned putative biological functions based on the sequence similarities.

Genome comparisons revealed that the newly isolated pelagiphages are all closely related to previously reported pelagiphages HTVC019P and HTVC011P, with sequence similarities and overall conservation of genome architectures. These HTVC019P-related pelagiphages belong to a well-characterized *Autographivirinae* subfamily within the *Podoviridae* family. Most pelagiphages fall within the criteria of >40% of the shared genes, with the exception of HTVC011P and HTVC025P, which share 32-46% genes with other pelagiphages. Considering their genome synteny and biological features, we group all HTVC019P-related pelagiphages into a new bacteriophage genus that is designated the *HTVC019Pvirus.*

tRNA genes were identified from five *HTVC019Pvirus* pelagiphage genomes (Fig. 1). HTVC011P and HTVC025P both encode a tRNA-Leu (TAG) gene downstream of the integrase, upstream of the RNA polymerase. While the tRNA-leu (TAA) encoded by HTVC200P and HTVC119P are located upstream of the integrase. HTVC019P encodes a tRNA-Cys (GCA) located between lysozyme and DNA primase (Fig. 1).

**Fig 1.**
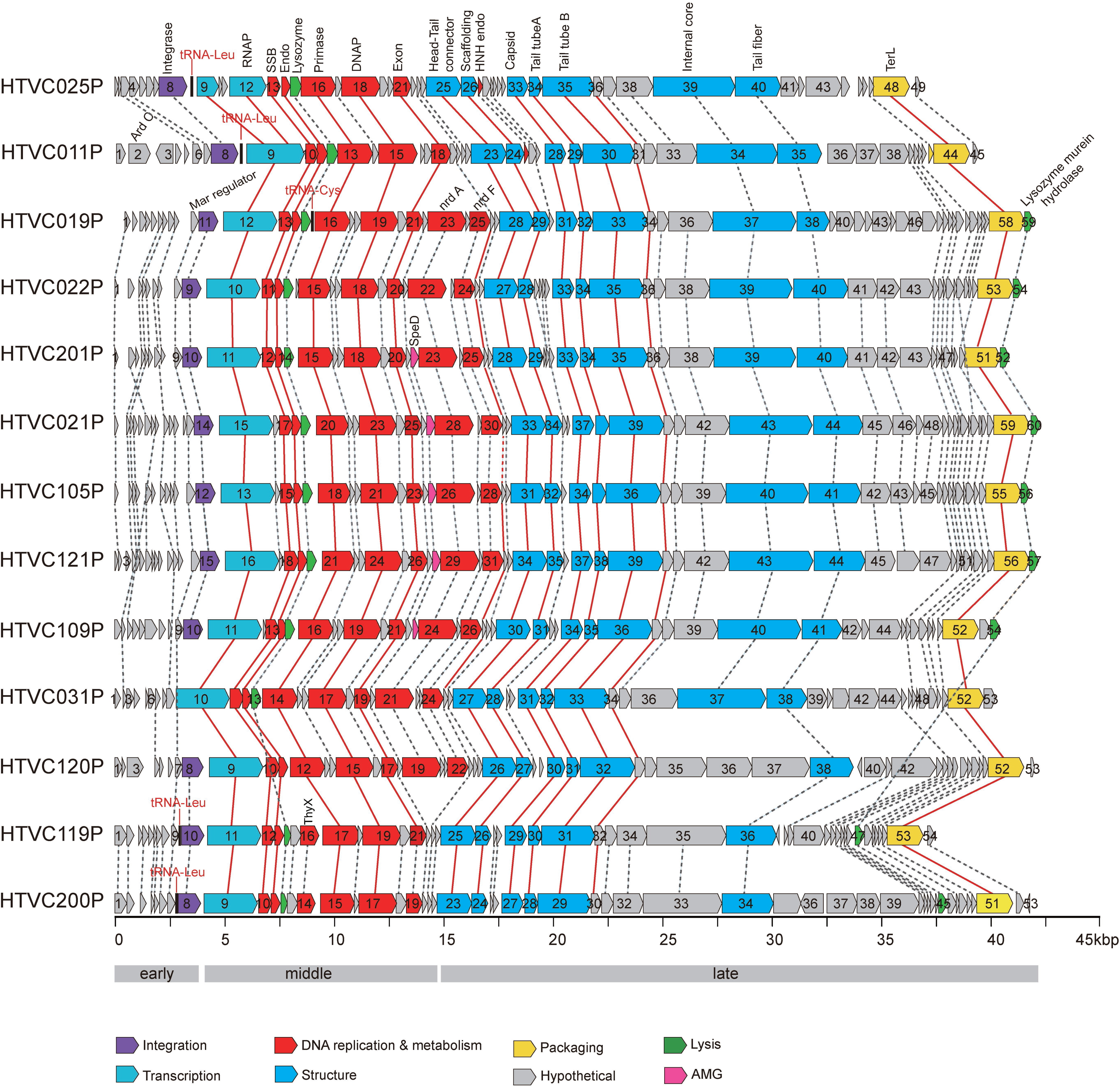
Genome organization and comparison of 13 *HTVC019Pvirus* pelagiphages. ORFs are depicted by leftward or rightward oriented arrows according to their transcription direction. The number of each ORF is shown within each arrow. ORFs are color-coded according to putative biological function. Shared ORFs are connected by dash lines and core genes are connected by red lines. tRNAs are shown in red. Abbreviation: MarR, MarR family transcriptional regulator; int, integrase; RNAP, RNA polymerase; SSB, single-stranded DNA binding protein; endo, endonuclease; DNAP, DNA polymerase; exon, exonuclease; SpeD, S-adenosylmethionine decarboxylase proenzyme; nrdA, ribonucleotide-diphosphate reductase alpha subunit; nrdF, ribonucleotide-diphosphate reductase beta subunit; TerL, terminase, large subunit.

HTVC021P and HTVC105P are similar (90% average nucleotide identity over 70% of their genomes), with most genome variation located in the module of genes encoding structural features. They infect different hosts and were isolated from geographically distant sampling sites (Table 1). Similarly, HTVC019P and HTVC022P infecting HTCC1062 are also similar (94% average nucleotide identity over 69% of their genomes) and were isolated from geographically distant sampling sites (Table 1). These findings suggest that pelagiphages were transferred to different ocean areas and some diverged to infect different hosts recently.

### Core and pan-genomes

Given the 13 complete *HTVC019Pvirus* pelagiphage genomes, we performed a core and pan genome comparative analysis. The *HTVC019Pvirus* pan-genome contains a total of 234 predicted protein clusters. As expected, the pan-genome accumulation curve did not saturate (Fig. 2), suggesting the need for more extensive investigation of *HTVC019Pvirus* pelagiphage pan genomes. A total of 15 core genes were found common among all *HTVC019Pvirus* pelagiphages, with extensive conservation of syteny (Fig. 1 and Fig. 2). These core genes possess the functions essential for the phage life cycle, including phage RNA polymerase catalyzed transcription, DNA metabolism and replication, cell lysis, phage structure and DNA maturation. The core genome of *HTVC019Pvirus* pelagiphages and P60-like cyanophage genomes share 12 genes (23) (see Table S1 in the supplemental material), suggesting that functional core gene composition of *HTVC019Pvirus* pelagiphages is similar to P60-like cyanophages.

**Fig 2.**
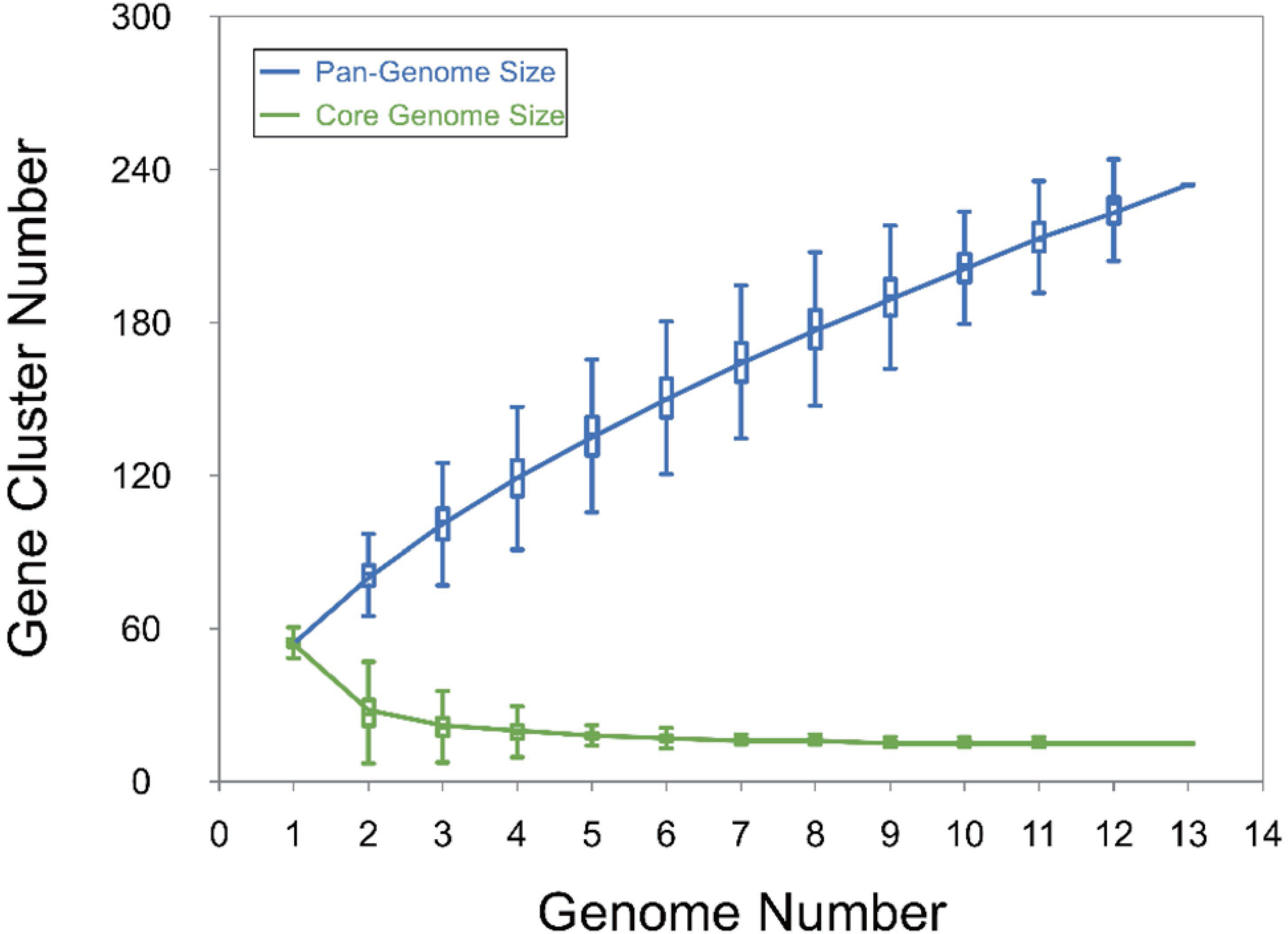
Pan- and core-genomes of *HTVC019virus* pelagiphages. Number of total genes in the core (green) and pan (blue) genomes as a function of the number of genomes included in the analysis.

### Phylogenomic analysis

To determine the phylogenetic relationships within the *HTVC019Pvirus* pelagiphages, a whole-genome phylogenies were inferred from 12 concatenated core genes. Twenty-six *HTVC019Pvirus* pelagiphages, including 13 from MedDCM fosmid library, were included in the analysis. The results show that *HTVC019Pvirus* pelagiphage isolates are separated into three major groups (group I-III) and an extra group composed of only MedDCM sequences (Fig. 3A). This result is consistent with gene content analysis (Fig. 3B), in which >50% genes are shared within groups, and 32-53% genes are shared between groups.

**Fig 3.**
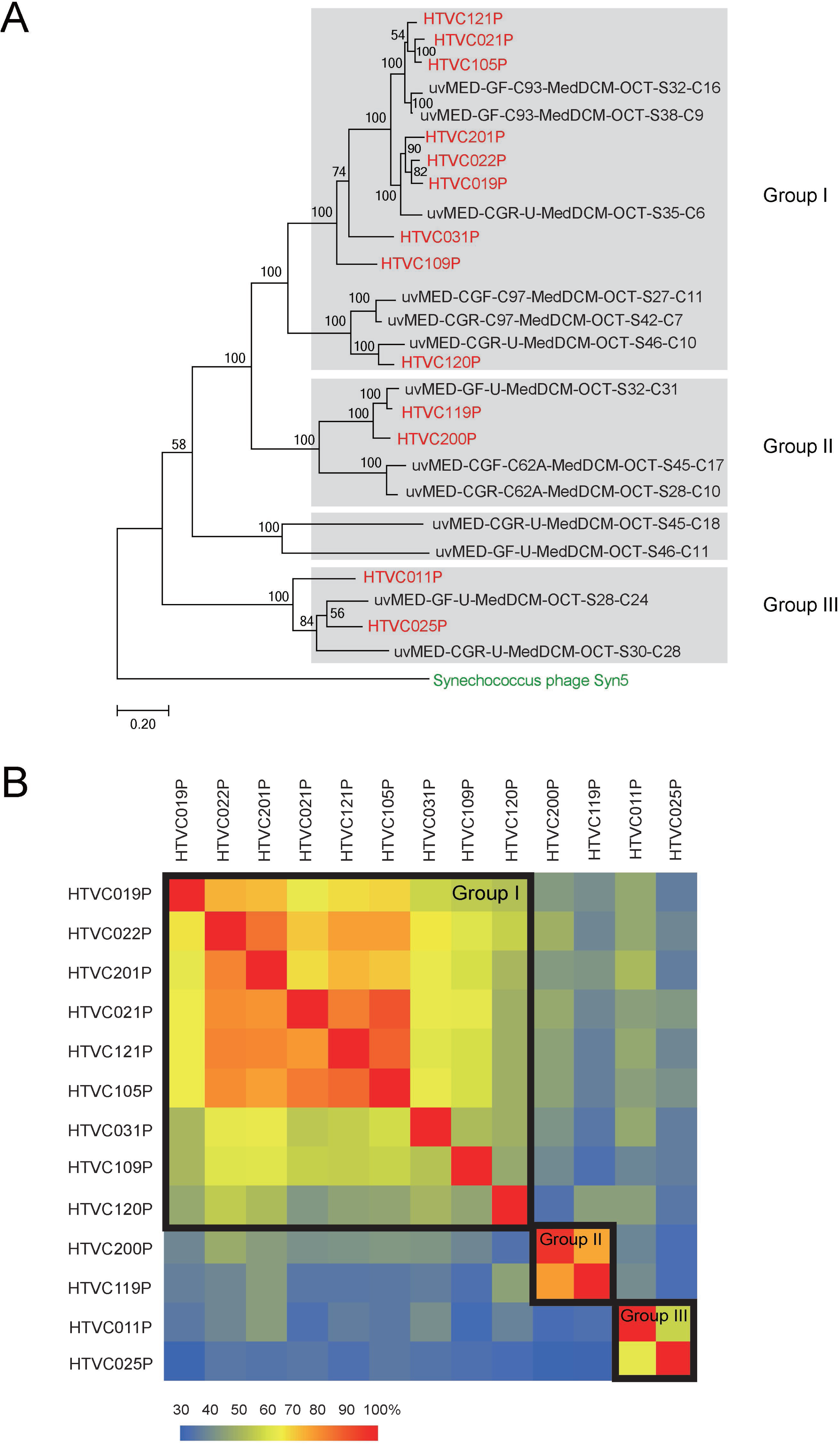
**A**. Maximum-likelihood phylogenetic trees of conserved phage core proteins. *Synechococcus* phage Syn5 shown in green is outgroup, pelagiphages in this study are shown in red, and the pelagiphage contigs from Mediterranean DCM (MedDCM) fosmid library were shown in black. The scale bar represents 0.2 fixed substitution per amino acid position. Bootstrap = 500. Only bootstrap values >50 are shown. **B**. The heatmap shows the percentage of shared genes between 13 pelagiphages. Phages in the same subgroups are boxed.

### Genome structure

Similar to other *Autographivirinae* phage genomes, *HTVC019Pvirus* pelagiphage genomes can be divided roughly into early, middle and late regions (Fig. 1). The early region is composed mostly of a suit of small proteins with unknown functions. Interestingly, all but one *HTVC019Pvirus* pelagiphage contain an integrase gene located upstream of the RNA polymerase. The integrase was not identified in the HTVC031P genome, which suggests that this phage is a putative obligate lytic phage. Homologs of phage T7 antirestriction protein Ocr, which is an inhibitor of the type I restriction-modification enzyme, were not found in these *HTVC019Pvirus* pelagiphage genomes. Instead, HTVC011P codes for a putative antirestriction protein (ArdC, PF08401.10) which shares 32% amino acid identity with an ArdC encoded by the Inc W plasmid pSa. The pSa ArdC protein can specifically repress type I restriction enzymes and it is also able to protect single-stranded DNA against endonuclease activity (25). It is possible that this phage encoded ArdC antirestriction protein can help phage DNA overcome the host restriction system. The middle genome region codes for the genes mainly involved in RNA transcription and DNA metabolism. All *HTVC019Pvirus* pelagiphages code for an RNA polymerase. A phage encoded DNA-dependent RNA polymerase responsible for most phage gene transcription is the hallmark of the *Autographivirinae* subfamily. All *HTVC019Pvirus* pelagiphages possess the typical T7-like DNA replication system including single-stranded DNA-binding protein (SSB), DNA polymerase, endonuclease I, DNA primase and exonuclease. All but one of *HTVC019Pvirus* pelagiphage possess a lysozyme gene sharing identity with the T7 lysozyme (36-41% amino acid identity). Phage lysozyme cuts the amide bonds in the bacterial cell wall, which is essential for cell lysis. T7-encoded lysozyme is a bifunctional enzyme, which possesses both amidase activity and transcription inhibition activity, by binding to T7 RNA polymerase (26, 27). Additionally, nine *HTVC019Pvirus* pelagiphages contain an extra gene in the late genome region, lysozyme murein hydrolase, that may be involved in cell lysis (Fig. 1). These lysozyme murein hydrolase genes share identity with hydrolase from T4-like myoviruses, which hydrolyses the beta-1,4-glycosidic bond between N-acetylmuramic acid (MurNAc) and N-acetylglucosamine (GlcNAc) in peptidoglycan heteropolymers of cell walls (28).

The late genome region codes for the genes involved in phage particles assembly, DNA packaging and cell lysis. All the *HTVC019Pvirus* pelagiphages encode a subset of virion structural proteins (Fig. 1). In addition, they all encode a terminase large subunit (TerL), sharing sequence identity with the T7 counterpart. T7 DNA maturases are involved in cutting DNA monomer from the concatemers and packaging DNA into phage heads. However, the small terminase subunit was not identified.

### Auxiliary metabolic genes (AMGs)

Like other bacteriophage genomes, *HTVC019Pvirus* pelagiphage genomes display mosaicism (29) and contain some genes potentially arising from horizontal genetic exchange. Some genes we identified in the *HTVC019Pvirus* genus pan genome may be auxiliary metabolic genes (AMGs) presumably acquired from their hosts (30). AMGs sometimes are thought to reinforce phage adaptation and fitness by modulating host metabolism in low-nutrient marine environments.

Genes encoding the two subunits of class I ribonucleotide reductases (*nrdA* and *nrdF)* were identified from all *HTVC019Pvirus* pelagiphages except for HTVC011P, HTVC031P and HTVC119P. Ribonucleotide reductases (RNRs) are common AMGs found in sequenced marine podoviruse genomes, including *Autograhivirinae* cyanophages, Roseophage SIO1 and N4-like roseophages (20, 21, 31-33). RNRs catalyze the formation of deoxyribonucleotides from ribonucleotides, providing the precursors required for DNA synthesis and repair (34). It was suggested that phages gained RNR genes from bacteria in order to obtain sufficient free nucleotides for DNA synthesis in phosphorus-limited marine environments (20, 33). Two pelagiphages, HTVC119P and HTVC200P, code for a thymidylate synthetase (thyX), which is also involved in nucleic acid synthesis and metabolism.

Interestingly, a gene encoding S-adenosylmethionine decarboxylase proenzyme (*speD*) (PF02675.14, E.C.4.1.1.50) was found in five *HTVC019Pvirus* pelagiphage genomes. *speD* is involved in the decarboxylation of S-adenosylmethionine to S-adenosylmethioninamine, and therefore is critical for biosynthesis of the polyamines spermine and spermidine by providing the propylamine donor (35). In bacteriophage T4, polyamines are involved in the DNA charge balance during phage genome packaging process (36). Polyamines is also an important substrate for the growth of SAR11 (16). It is possible that *speD* benefits both host and phage during infection. *speD* was also identified from marine T4-like cyanomyophage genomes and a unique polar freshwater cyanophage genome (37-40).

### Phage integrase genes and identification of integration sites

As described above, 12 *HTVC019Pvirus* pelagiphages harbor an integrase gene upstream of the RNA polymerase. Integrase genes usually occur in temperate phage genomes and are responsible for site-specific recombination between *attB* and *attP*. Based on the sequence homology and catalytic residues, all *HTVC019Pvirus* pelagiphage integrases belong to the tyrosine integrase family. Sequence alignment shows that most of the integrases contain the catalytic residue tetrad Arg-His-Arg-Tyr (R-H-R-Y) in the C-terminal catalytic domain which is needed for DNA cleavage and joining (41, 42) (Fig. S1). However, in HTVC025P integrase, Tyr substitutes for His (R-Y-R-Y). In HTVC011P integrase, the His site and the second Arg site were substituted with Tyr and Lys residues, respectively (R-Y-K-Y)

To investigate *HTVC019Pvirus* pelagiphage integration to form lysogenized host cells, several strategies were used to identify the phage integration sites: (i) First, phage integration sites were discovered by high-throughput sequencing analysis. As a result of phage integration, the integrated phage genome is inserted between left and right attachment sites, *attL* and *attR*. The sequences containing the hybrid *attL* and *attR* sites were identified from illumina sequencing data, allowing the identification of the *attB* and *attP* sites. (ii) Second, PCR assays were used to confirm the phage site-specific integration. PCR primers targeting the *attL* and *attR* sites were designed to amplify the left and right integration flanking regions from phage-infected host cultures (primer sets are indicated in Fig. 4 and listed in Table S3). PCR fragments of expected size were successfully amplified. Comparison of the PCR sequencing results showed the expected junction fragments. (iii) Metagenome searches were performed to detect the integration sites. GOS database searches were carried out using phage integrase and RNA polymerase sequences as queries, yielding fragments containing both pelagiphage and SAR11 homologues (discussed below).

**Fig 4.**
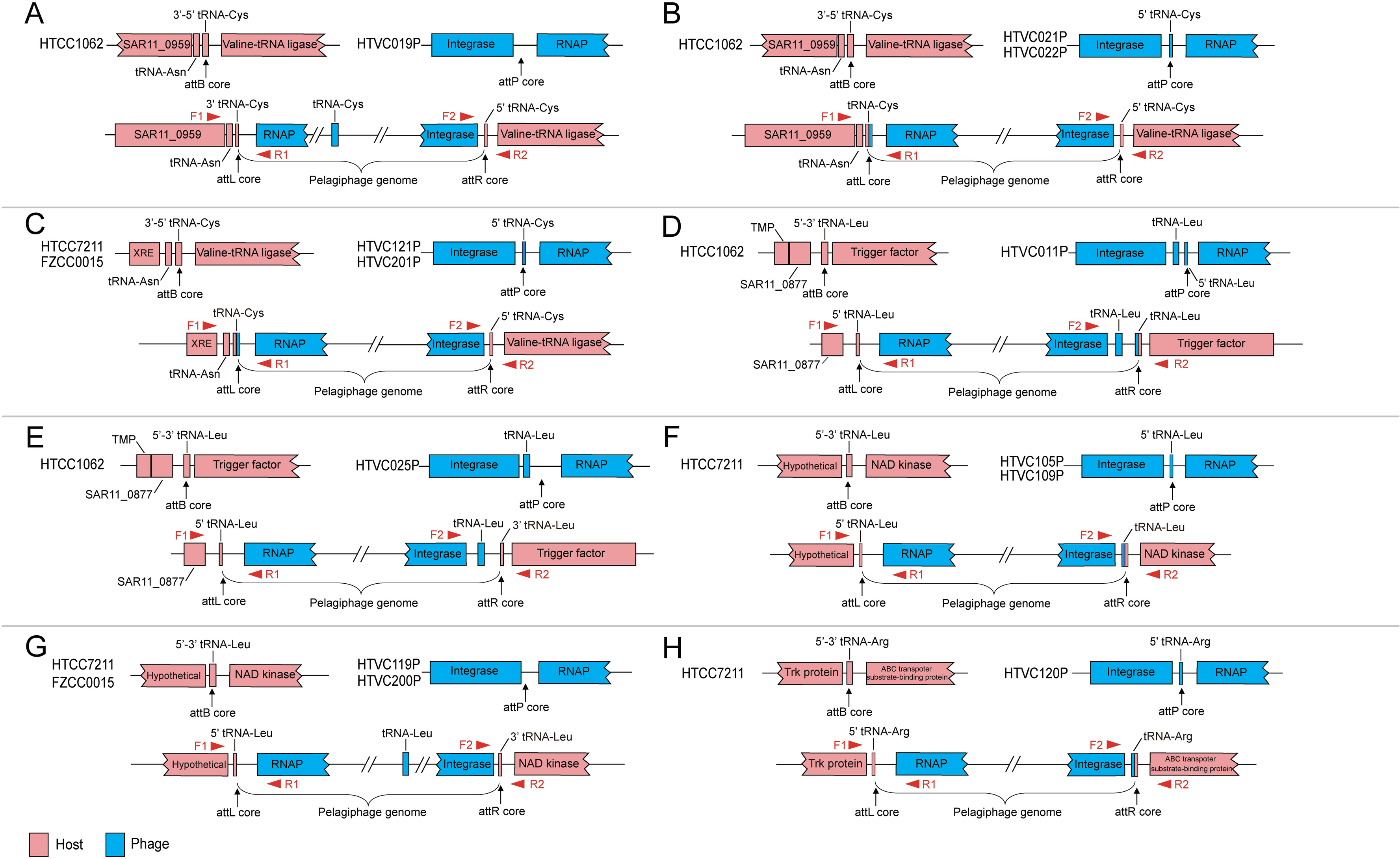
The genome organization around the *att*B and *att*P in the pelagiphage and host genomes. A. HTVC019P; B. HTVC021P and HTVC022P; C. HTVC121P and HTVC201P; D. HTVC011P; E. HTVC105P and HTVC109P; F. HTVC119P and HTVC200P; G. HTVC120P. The location of the core sequence regions within the integration sites (*att*B, *att*P, *att*L and *att*R) are indicated by the arrows. Host and phage genes are shown in pink and blue, respectively. PCR primers are indicated by triangles. See Supplementary Fig. S2 for sequences of all integration sites. Abbreviation: RNAP, RNA polymerase; TMP, Transmembrane protein; XRE, XRE family transcriptional regulator.

### Phage integration sites and core sequences

The genome organization around the integration sites are shown in Fig. 4 and summarized in Table S2. Our analysis revealed the integration sites of 12 *HTVC019Pvirus* pelagiphages. In these pelagiphages, all *attP* sites are located in the non-codeing region between integrase and RNA polymerase. While, in the SAR11 genomes, the *attB* sites are located within various tRNA genes. Bacterial tRNA genes or adjacent sequences of tRNA genes are known to be common integration sites for prophages (43, 44). Sequences comparisons revealed a common 11-46 bp ‘core sequence’ shared by all attachment sites, where the site-specific recombinations take place (Table S2 and Fig. S2).

The analysis of the phage-host junction fragments suggested that HTVC019P, HTVC021P and HTVC022P can all integrate into the HTCC1062 genome at a tRNA-Cys (GCA) site (Fig. 4A and 4B). The core sequence is located in the middle of the HTCC1062 tRNA-Cys gene while the *attP* core sequence lies downstream of the integrase gene in HTVC019P (Fig. 4A and Fig. S2). Notably, upon HTVC019P integration, the HTCC1062 tRNA-Cys gene was separated into two parts and is presumed to be non-functional (Fig. 4A). The tRNA-Cys gene located between lysozyme and DNA primase in HTVC019P is likely used instead (Fig. 4A). In the case of HTVC021P and HTVC022P, the identical core sequence is located in the vicinity of the 5’ end of the host tRNA-Cys gene (Fig. 4B and Fig. S2). Upon phage integration, the HTCC1062 tRNA-Cys gene was disrupted and complemented by the partial 5’ end of tRNA-Cys gene in phage. The reconstituted tRNA-Cys carries a base alteration at phage-derived 5’ end (indicated by arrow in Fig. S2).

Similarly, HTVC201P and HTVC121P are able to integrate at the tRNA-Cys (GCA) site in the FZ0015 and HTCC7211 genome, respectively (Fig. 4C). The core sequences of both phages overlap the 5’ end of the host tRNA-Cys gene (Fig. S2). Upon phage integration, owing to the identical core regions, both tRNA-Cys genes do not show any alteration (Fig. S2).

The integration site of HTVC011P and HTVC025P in the HTCC1062 genome is the tRNA-Leu (TAG) gene. The core sequences are both located in the middle of the host tRNA-Leu gene (Fig. S2). Note that, in HTVC011P, the core sequence lies downstream of the tRNA-Leu gene, where exist 5’ end of a tRNA-Leu gene (Fig. 4D). Upon phage integration, the host tRNA-Leu gene was separated into two parts and complemented by the partial tRNA-Leu gene in phage. Consequently, there exist two tRNA-Leu genes around the *attR* site after HTVC011P integration (Fig. 4D). In the case of HTVC025P, upon phage integration, the host tRNA-Leu gene was discrupted but could not be complemented, thus the tRNA-Leu gene in HTVC025P is likely used instead (Fig. 4E).

HTVC105P, HTVC109P and HTVC119P are all able to integrate at a tRNA-Leu (TAA) site in the HTCC7211 genome (Fig. 4F and 4G). In the case of HTVC105P, the core sequence overlap the 5’ end of the host tRNA-Leu gene (Fig. S2). Upon HTVC105P integration, the tRNA-Cys does not show any alteration. In HTVC109P and HTVC119P, the core sequences are located in the middle of the host tRNA-Leu gene (Fig. S2). Upon HTVC109P integration, the host tRNA-Leu gene was separated into two parts and complemented by the partial 5’ end of the tRNA-Leu in phage (Fig. 4F). Consequently, the reconstituted tRNA-Leu gene carries some alteration (deletion) at phage-derived 5’ end (indicated by arrow in Fig. S2). While upon HTVC119P integration, tRNA-Leu was also disrupted but could not be complemented by the phage sequence (Fig. 4F). The tRNA-Leu gene located upstream of the integrase in HTVC119P is likely used instead (Fig. 4G). Similar to HTVC119P, HTVC200P can integrate at the tRNA-Leu (TAA) gene in FZCC0015. After HTVC119P integration, tRNA-Leu was disrupted and the tRNA-Leu gene in HTVC200P is likely used instead (Fig. 4G)

HTVC120P targets the tRNA-Arg site in the HTCC7211 genome (Fig. 4H). The core sequence is in the middle of the host tRNA-Arg gene. Upon phage integration, the disrupted host tRNA-Arg gene was complemented by the partial 5’ end of the tRNA-Cys in phage, resulting in a reconstituted tRNA-Arg that carries two base alterations at phage-derived 5’ end (indicated by arrow in Fig. S2).

### GOS recruitment of hybrid sequences

We explored whether pelagiphage integration can also be detected in environmental datasets by using metagenome search strategy. After annotation and analysis, one sequence was found containing the homologues of the C-terminal end of the SAR11 transmembrane protein (TMP), 5’ end of a tRNA-Leu gene, and the N-terminal end of the pelagiphage RNA polymerase (Fig. 5A). This sequence indicates the *attL* site of pelagiphage integration. Five sequences were found containing homologues of the C-terminal end of the pelagiphage integrase, an intact tRNA-Leu gene, 3’ end of a tRNA-Leu gene, and the N-terminal of the SAR11 trigger gene, indicating the *attR* sites (Fig. 5B). Additionally, a fragment containing homologues of the C-terminal of the pelagiphage integrase, an intact tRNA-Cys gene, and the N-terminal end of the SAR11 Valine-tRNA ligase was retrieved (Fig. 5C). The discovery of these phage-host junctions suggests that pelagiphages can integrate into the SAR11 genomes at the tRNA-Leu and tRNA-Cys sites *in situ*.

**Fig 5.**
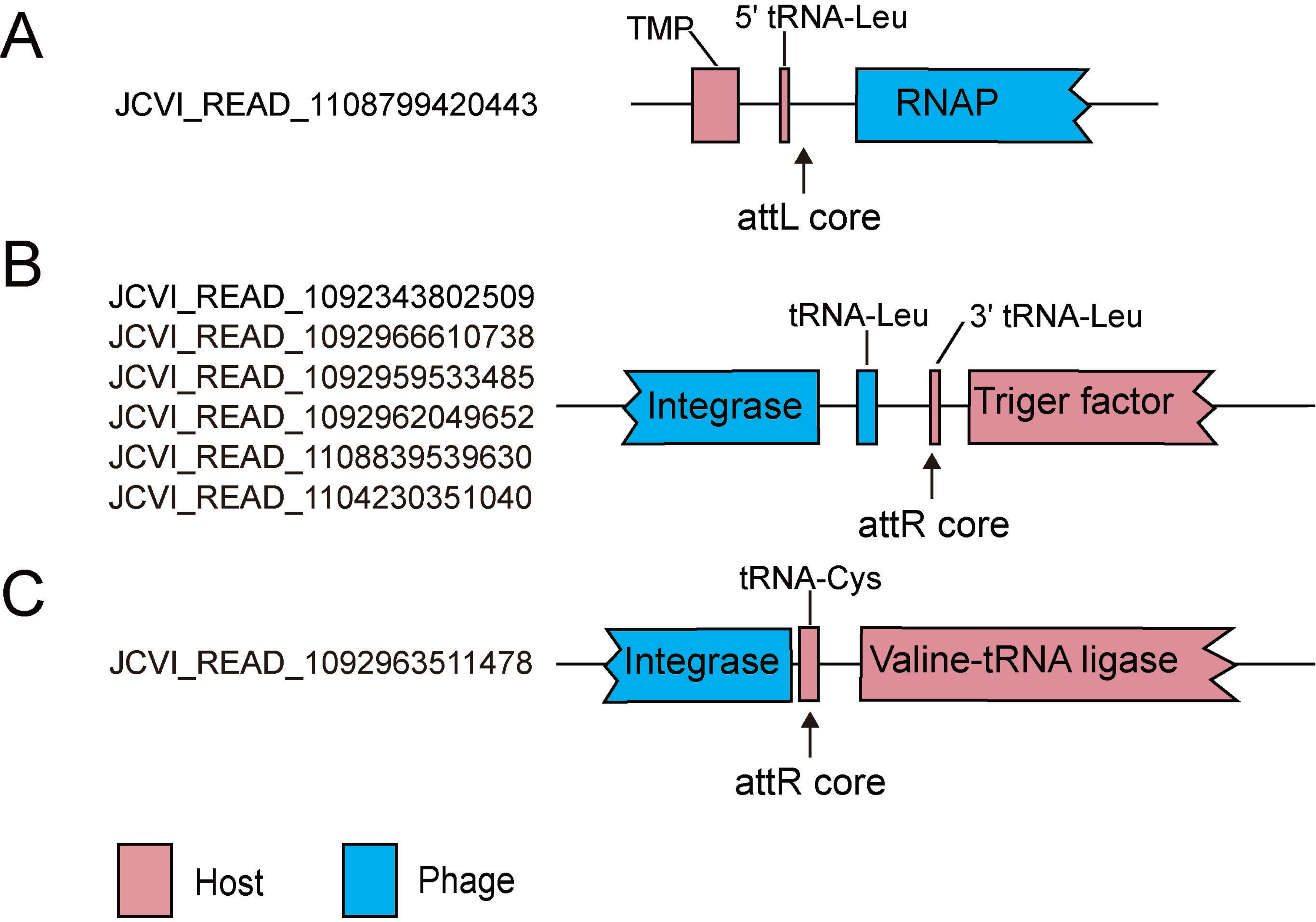
Hybrid fragments retrieved from GOS database verifying the occurrence of pelgiphage integration in situ. **A**. sequence around *attL* site of pelagiphage integration into the tRNA-Leu. **B**. sequences around *attR* site of pelagiphage integration into the tRNA-Leu. **C**. sequence around attR site of pelagiphage integration into the tRNA-Cys. Host and phage genes are shown in pink and blue, respectively. The location of the core sequence regions within the integration sites are indicated by the arrows.

The importance of prophages in marine systems has been recognized for a decade. Researcher found that about half of marine bacterial genomes contain prophage-like elements (45). However, no prophage-like elements have been observed in sequenced SAR11 genomes. In our study, phage integration mechanisms were demonstrated with phage isolates growing in culture and metagenomic searches. Given the prevalence of *Autographivirinae* pelagiphages in the ocean and our obsvervation of lysogeny, our results suggest that *Autographivirinae* prophage are an important prophage type in the ocean.

Phage lysogeny can provide a mutually beneficial relationship to bacteria and phages (1, 45, 46). Lysogeny can benefit phages under circumstances where selection favors propagation of the lysogen. Lysogeny also provides bacteria with multiple benefits, including providing genetic content and new phenotypic traits, providing immunity to other related phage infections and increasing inter-strain genetic variation (45, 47).

In the well-studied phage lambda-*E.coli* model system, it was demonstrated that lambda CI-Cro bi-stable switch controls the decision between the lytic and lysogenic pathways (48). Unfavorable environmental conditions, including nutrient deficiency, lower temperature, and high multiplicity of infection (MOI), favor the establishment of lysogeny (49). In natural marine environments, it has been suggested that low nutrients, low system productivity and slow bacterial growth rates favor the lysogenic strategy (59,50-53). Phosphate limitation has been proposed as principal trigger responsible for lytic-to-lysogeny switching (reviewed by Wilson and Mann 1997) (54). In contrast, the recently proposed a “Piggyback-the-Winner (PtW)” hypothesis explains environmental trends in virus/bacteria ratios by postulating that the lysogeny strategy is preferred under conditions of high host densities and rapid host growth rate (2, 55).

The prevalence of pathways for lysogeny in *HTVC019Pvirus* pelagiphages remains a puzzle. It is unclear what factors determine whether these pelagiphages enter the lysogenic or lytic cycle upon infection. Phage lysogen frequencies are also unknown. Further study will be required to elucidate the underlying genetic mechanisms that control the pelagiphage decision between the lytic and lysogenic pathways.

Despite the advantages of lysogeny to bacteria, integrated phage genomes could pose extra metabolic and fitness burden to bacteria. SAR11 is known to have small cell size, streamlined genome structure with few pseudogenes and minimized intergenic spacers (56-59). Genome streamlining is critical to the success of SAR11 in nutrient-limited marine environments (59). Neither plasmid nor prophage-like elements were found in any sequenced SAR11 genomes, which implies the genome streamlining theory. However, on the contrary, our study revealed that prophage integration, which will increase the resource requirement for the bacterial replication, is present in SAR11 genomes. It is plausible that the benefits of carrying a prophage which account for about ~2-3% of SAR11 genome compensate the energetic cost of replicating prophages. The lack of SAR11 isolates containing prophage may be due the slow growth rate and low density of lysogenic SAR11 cells, thus were difficulty to be isolated from the ocean by using high throughput culturing method. Considering that lysogenized SAR11 cells have not yet been obtained, it is still unknown whether the integrated pelagiphages can maintain a stable symbiotic relationship with their host. According to a recent analysis of 2110 bacterial genomes, prophages are rare in bacteria with slow growth rates and small genome sizes (60). It is possible that prophage is not able to stably integrate during the infection, the prophage-SAR11 genome coexistence is temporary in order to keep host genome streamlined. Recently, a “lyso-lysis” phenomenon was observed in bacteriophage lambda, where phage integration was followed by a lytic life cycle (61) and the frequency of lyso-lysis increases with the number of infecting phages. Even this bacteria-phage genome coexistence is likely temporary, still provides a window for evolution of mutually benefical functions (62).

Considering that most yet-known *Autographivirinae* prophage structures are found in non-marine bacterial genomes, the possession of lysogenic capability apparently is not a typical feature of marine *Autographivirinae* phages. The evolutionary origins of integrase genes in the *Autographivirinae* phage genomes are still unclear. It is possible that lytic phages gained the integrase genes through lateral gene transfers. It is also possible that integrase genes were lost in some phage genomes during their evolutionary histories, resulting in an exclusively lytic life cycle.

### Conclusions

The study of new *HTVC019Pvirus* pelagiphages provided new insights into the life cycles, diversity, genomic adaptations and evolution of this abundant virus group. We identified phage integration sites and demonstrated that most *HTVC019Pvirus* pelagiphages are capable of lysogenizing their hosts. The capacity of *HTVC019Pvirus* pelagiphages for either lysing their hosts or integrating into the host genome suggests they impact on SAR11 populations by a variety of mechanisms, including mortality, genetic exchange, and possibly viral-induced immunity. Details of *HTVC019Pvirus* integration and excision and the mechanism that controls the decision between the lytic and lysogenic pathways will be an avenue of future research. The prevalence of *HTVC019Pvirus* pelagiphages in marine systems makes them potentially important as an experimental model for understanding phage strategies and evaluating conceptual models such as the viral shunt, Kill-the-Winner, Piggyback-the-Winner, and King-of-the-Mountain.

## MATERIALS AND METHODS

### Host cells

*Pelagibacter* sp. str. HTCC7211 and *Pelagibacter* HTCC1062 were grown in artificial seawater based medium, amended with FeCl_3_, pyruvate, glycine, methionine and vitamins (63). Strain FZCC0015 was isolated on 13th May, 2017 from the coast of Pingtang island in China (lat.’ N25°26′, long. E119°47′) by using dilution-to-extinction method (64). *Pelagibacter* FZCC0015 has been fully sequenced (Accession number: CP031125). FZCC0015 was grown in a seawater based medium with excess vitamin and amended with NH_4_Cl, KH_2_PO_4_, FeCl_3_, pyruvate, glycine, and methionine.

### Source waters and phage isolation

Water samples were collected from a variety of ocean sampling stations, from surface to 150 m (Table 1). Water samples were filtered through 0.1 μm filters to obtain a bacteria-free fraction. The filtered samples were stored in the dark at 4°C until use. The pelagiphage isolation procedure has been described in detail previously (14). Briefly, seawater samples were inoculated with host cultures and monitored for the cell lysis by using a Guava easyCyte Flow Cytometer (MerckMillipore, Billerica, MA). When cell lysis was detected, the presence of phage particles was confirmed by epifluorescence microscopy (65). Purified phage clones were obtained by using the dilution-to-extinction method. The purity of pelagiphages was examined by genome sequencing.

### Cross-infection experiments

Cross-infection experiments were performed by using 13 pelagiphages infect against three SAR11 hosts. Exponentially growing cultures of SAR11 strains were incubated with each single pelagiphage at a phage-to-host ratio of ~20. Cell growth was monitored by flow cytometry and phage particles were enumerated by epifluorescence microscopy.

### Phage DNA preparation, genome sequencing and annotation

Four liters of host cultures were infected with each pelagiphage at a phage-to-host ratio of ~3. Following host lysis, cell debris was removed by centrifugation at 10 000 rpm for 60 min. Phage lysates were then concentrated using Pellicon 2 mini filter tangential flow filtration system with a 30-kDa (MerckMillipore, Bedford, MA). Concentrated phage lysates were further concentrated by Amicon Ultra Centrifugal Filters (30-kDa, MerckMillipore). Phage DNA was extracted using a formamide extraction method (66) and sequenced by using Illumina paired-end HiSeq 2500 sequencing approach by Mega genomics Technology Co., Ltd (Beijing, China). The reads were quality-filtered, trimmed and de novo assembled using CLC Genomic Workbench 11.0.1 (QIAGEN, Hilden, Germany). The GenBank accession numbers assigned to the complete pelagiphages genomes are listed in Table 1.

Putative open reading frames (ORFs) longer than 120 bp were identified from phage genomes using a combination of Glimmer 3.0 (67), RAST server (68), GeneMark (69), and manual inspection. Putative biological functions were assigned to translated ORFs using BLASTP against the NCBI non-redundant (nr) and NCBI-refseq databases. In this study, genes with >30% amino acid identity, >50% alignment coverage of the shortest protein, and an E-value cutoff <1E-3 were considered to be homologs. PFAM and HHpred were also employed to identify the protein families. The tRNAs were identified using tRNAscan-SE (70).

### Phylogenomic analysis

12 core genes were selected for phylogenomic analysis (Table S1). Individual alignments for each of the core genes were constructed with MUSCLE (71) and edited with Gblocks (72). Alignment of the concatenated core genes was evaluated for optimal amino acid substitution model using ProtTest (73), and run with RAxML v8 (74).

### Pan genome and core genome

The protein sequences of 13 pelagiphages were clusters into orthologous groups using OrthoMCLv2.0 with an inflation index of 1.5 (75, 76). OrthoMCL used the following BLASTP parameters: E-value cut-off of 1e-5, 50% minimum aligned coverage and 30% minimum amino acid identity. Gene accumulation curves were generated using R script for OrthoMCL results.

### Determination of pelagiphage integration sites

Exponentially-growing SAR11 bacteria cells were infected separately by single pelagiphage. When cell lysis was observed, phage-infected cells were harvested by centrifugation at 20 000 rpm for 30 min. DNA were extracted from the host cells using DNeasy Blood & Tissue kit (Qiagen, Germantown, MD). DNA samples were sequenced by using Illumina paired-end HiSeq 2500 sequencing approache by Novogene Technology Co., Ltd (Beijing, China). The reads were quality-filtered, trimmed and mapped to pelagiphage genomes using CLC Genomic Workbench 11.0.1. Sequences mapped to phage genomes was manually inspected to find the phage-host hybrid sequences. The resulting sequences were analyzed to identify integration sites and their locations. PCR primer sets were designed based on the predicted *att*L and *att*R sites. The location of each primer set was indicated in Fig. 4, and the primer sequences were provided in Table S3. PCR was performed in a 50 μl volume containing 1×PCR buffer, 2 mM MgSO_4_, 0.2 mM of each deoxynucleoside triphosphate, 0.2 μM of each primer, and 1 U Taq enzyme. DNA extracted from phage infected host cells was used as PCR template. The PCR program for all reactions included an initial denaturing step at 95°C for 3 min, followed by 35 cycles of 95°C for 1 min, annealing at 55°C for 30 s, and extension at 72°C for 1 min, followed by a final extension step at 72°C for 10 min.

### Metagenomic search for pelagiphage integration sites

Given that the adjacent locations of integrase and integration site are found in most temperate phage genomes, amino acid sequences of integrase and RNA polymerase from pelagiphages were used as queries to search against Global Ocean Survey (GOS) metagenomic database using tBLASTN, with e-value threshold of 1E-3. The resulting fragments containing homologs of phage integrase or RNAP were searched against the IMG database and a SAR11 genome dataset using BLASTx. Only the fragments also containing sequences best hit to SAR11 genes were retained for further analysis.

## SUPPLEMENTAL MATERIAL

**Fig S1,** TIF file

**Fig S2,** TIF file

**Table S1,** DOCX file

**Table S2,** DOCX file

**Table S3,** DOCX file

## ACKNOWLEDGMENTS

The study was supported by NSFC grant 41706173 and GBMF607.01. We thank Ben Knowles for the useful suggestions.

## Supplementary Figure Legends

**Fig S1** Mutiple protein sequence alignment of integrase catalytic domains. The conserved residues (R-H-R-Y) are highlighted in red.

**Fig S2** Alignment of DNA sequences around attachment sites. The bacterial chromosomal sequences, pelagiphage sequences, and identical core sequences are in blue, black and red, respectively. The tRNA genes found in the integration sites are boxed. The changed positions in the tRNAs are indicated by arrows.

## REFERENCES

1. Wommack KE, Colwell RR. 2000. Virioplankton: Viruses in aquatic ecosystems. Microbiol Mol Biol Rev 64:69–11 https://doi.org/10.1128/MMBR.64.1.69-114.2000.

2. Knowles B, Silveira CB, Bailey BA, Barott K, Cantu VA, Cobián-Güemes AG, Coutinho FH, Dinsdale EA, Felts B, Furby KA, George EE, Green KT, Gregoracci GB, Haas AF, Haggerty JM, Hester ER, Hisakawa N, Kelly LW, Lim YW, Little M, Luque A, McDole-Somera T, McNair K, de Oliveira LS, Quistad SD, Robinett NL, Sala E, Salamon P, Sanchez SE, Sandin S, Silva GG, Smith J, Sullivan C, Thompson C, Vermeij MJ, Youle M, Young C, Zgliczynski B, Brainard R, Edwards RA, Nulton J, Thompson F, Rohwer F. 2016. Lytic to temperate switching of viral communities. Nature 531:466–470. https://doi.org/10.1038/nature17193.

3. Wigington CH, Sonderegger D, Brussaard CPD, Buchan A, Finke JF, Fuhrman JA, Lennon JT, Middelboe M, Suttle CA, Stock C, Wilson WH, Wommack KE, Wilhelm SW, Weitz JS. 2016. Re-examination of the relationship between marine virus and microbial cell abundances. Nat Microbiol 1:15024. https://doi.org/10.1038/nmicrobiol. 2015.24.

4. Bergh Ø, Børsheim KY, Bratbak G, Heldal M. 1989. High abundances of viruses found in aquatic environments. Nature 340:467–468. https://doi.org/10.1038/340467a0.

5. Fuhrman JA. 1999. Marine viruses and their biogeochemical and ecological effects. Nature 399:541–548. https://doi.org/10.1038/21119.

6. Suttle CA. 2005. Viruses in the sea. Nature 437:356–361. https://doi.org/10.1038/nature04160.

7. Suttle CA. 2007. Marine viruses-major players in the global ecosystem. Nature Rev Microbiol 5:801–812. https://doi.org/10.1038/nrmicro1750.

8. Winter C, Bouvier T, Weinbauer MG, Thingstad TF. 2010. Trade-offs between competition and defense specialists among unicellular planktonic organisms: the “killing the winner” hypothesis revisited. Microbiol Mol Biol Rev 74:42–57. https://doi.org/10.1128/MMBR.00034-09

9. Thingstad TF, Lignell R. 1997. Theoretical models for the control of bacterial growth rate, abundance, diversity and carbon demand. Aqua Microb Ecol 13:19–27. https://doi.org/10.3354/ame013019

10. Thingstad TF. 2000. Elements of a theory for the mechanisms controlling abundance, diversity, and biogeochemical role of lytic bacterial viruses in aquatic systems. Limnol Oceanogr 45:1320–1328. https://doi.org/10.4319/lo. 2000.45.6.1320.

11. Lindell D, Sullivan MB, Johnson ZI, Tolonen AC, Rohwer F, Chisholm SW. 2004. Transfer of photosynthesis genes to and from Prochlorococcus viruses. Proc Natl Acad Sci U S A 101:11013–11018. https://doi.org/10.1073/pnas.0401526101.

12. Avrani S, Wurtzel O, Sharon I, Sorek R, Lindell D. 2011. Genomic island variability facilitates Prochlorococcus–virus coexistence. Nature 474:604–608. https://doi.org/10.1038/nature10172.

13. Marston MF, Pierciey FJ Jr, Shepard A, Gearin G, Qi J, Yandava C, Schuster SC, Henn MR, Martiny JB. 2012. Rapid diversification of coevolving marine Synechococcus and a virus. Proc Natl Acad Sci U S A 109:4544–4549. https://doi.org/10.1073/pnas.1120310109.

14. Zhao Y, Temperton B, Thrash JC, Schwalbach MS, Vergin KL, Landry ZC, Ellisman M, Deerinck T, Sullivan MB, Giovannoni SJ. 2013. Abundant SAR11 viruses in the ocean. Nature 494:357–360. https://doi.org/10.1038/nature11921.

15. Morris RM, Rappe MS, Connon SA, Vergin KL, Siebold WA, Carlson CA, Giovannoni SJ. 2002. SAR11 clade dominates ocean surface bacterioplankton communities. Nature 420: 806–810. https://doi.org/10.1038/nature01240.

16. Giovannoni SJ. 2017. SAR11 Bacteria: The most abundant plankton in the oceans. Ann Rev Mar Sci 9:231–255. https://doi.org/10.1146/annurev-marine-010814-015934.

17. Lavigne R, Seto D, Mahadevan P, Ackermann HW, Kropinski AM. 2008. Unifying classical and molecular taxonomic classification: analysis of the Podoviridae using BLASTP–based tools. Res Microbiol 159:406–414. https://doi.org/10.1016/j.resmic.2008.03.005.

18. Groth AC, Calos MP. 2004. Phage integrases: biology and applications. J Mol Biol 335:667–678. https://doi.org/10.1016/j.jmb.2003.09.082.

19. Nelson KE, Weinel C, Paulsen IT, Dodson RJ, Hilbert H, Martins dos Santos VA, et al. 2002. Complete genome sequence and comparative analysis of the metabolically versatile Pseudomonas putida KT24Environ Microbiol 4:799–80. https://doi.org/10.1046/j.1462-2920.2002.00366.x.

20. Sullivan MB, Coleman ML, Weigele P, Rohwer F, Chisholm SW. 2005. Three Prochlorococcus cyanophage genomes: signature features and ecological interpretations. PLoS Biol 3:e144. https://doi.org/10.1371/journal.pbio.0030144.

21. Pope WH, Weigele PR, Chang J, Pedulla ML, Ford ME, Houtz JM, Jiang W, Chiu W, Hatfull GF, Hendrix RW, King J. 2007. Genome sequence, structural proteins, and capsid organization of the cyanophage Syn5: a “horned” bacteriophage of marine synechococcus. J Mol Biol 368:966–981. https://doi.org/10.1016/j.jmb.2007.02.046.

22. Labrie SJ, Frois-Moniz K, Osburne MS, Kelly L, Roggensack SE, Sullivan MB, Gearin G, Zeng Q, Fitzgerald M, Henn MR, Chisholm SW. 2013. Genomes of marine cyanopodoviruses reveal multiple origins of diversity. Environ Microbiol 15:1356–1376. https://doi.org/10.1111/1462-2920.12053.

23. Huang S, Zhang S, Jiao N, Chen F. 2015. Comparative genomic and phylogenomic analyses reveal a conserved core genome shared by estuarine and oceanic cyanopodoviruses. PLoS One 10:e0142962. https://doi.org/10.1371/journal.pone.0142962.

24. Mizuno CM, Rodriguez-Valera F, Kimes NE, Ghai R. 2013. Expanding the marine virosphere using metagenomics. PLoS Genet 9:e1003987. https://doi.org/10.1371/journal.pgen.1003987.

25. Belogurov AA, Delver EP, Agafonova OV, Belogurova NG, Lee LY, Kado CI. 2000. Antirestriction protein Ard (Type C) encoded by IncW plasmid pSa has a high similarity to the “protein transport” domain of TraC1 primase of promiscuous plasmid RP4. J Mol Biol 296:969–977. https://doi.org/10.1006/jmbi.1999.3493.

26. Huang J, Villemain J, Padilla R, Sousa R. 1999. Mechanisms by which T7 lysozyme specifically regulates T7 RNA polymerase during different phases of transcription. J Mol Biol 293:457–475. https://doi.org/10.1006/jmbi.1999.3135.

27. Zhang X, Studier FW. 2004. Multiple roles of T7 RNA polymerase and T7 lysozyme during bacteriophage T7 infection. J Mol Biol 340:707–730. https://doi.org/10.1016/j.jmb.2004.05.006.

28. Jollès P, Jollès J. 1984. What’s new in lysozyme research? Mol Cell Biochem 63:165–189.

29. Hendrix RW, Smith MC, Burns RN, Ford ME, Hatfull GF. 1999. Evolutionary relationships among diverse bacteriophages and prophages: all the world’s a phage. Proc Natl Acad Sci U S A 96:2192–2197. https://doi.org/10.1073/pnas.96.5.2192.

30. Breitbart M, Thompson LR, Suttle CA, Sullivan MB. 2007. Exploring the vast diversity of marine viruses. Oceanogr 20:135–139. https://doi.org/10.5670/oceanog.2007.58.

31. Zhao Y, Wang K, Jiao N, Chen F. 2009. Genome sequences of two novel phages infecting marine roseobacters. Environ Microbiol 11:2055–2064. https://doi.org/10.1111/j.1462-2920.2009.01927.x.

32. Rohwer F, Segall A, Steward GF, Seguritan V, Breitbart M, Wolven F. Azam F. 2000. The complete genomic sequence of the marine phage Roseophage SIO1 shares homology with nonmarine phages. Limnol Oceanogr 45:408–418. https://doi.org/10.4319/lo.2000.45.2.0408.

33. Chen F, Lu J. 2002. Genomic sequence and evolution of marine cyanophage P60: a new Insight on lytic and lysogenic Phages. Appl Environ Microbiol 68:2589–2594. https://doi.org/10.1128/AEM.68.5.2589-2594.2002.

34. Nordlund P, Reichard P. 2006. Ribonucleotide reductases. Annu Rev Biochem 75:681–706. https://doi.org/10.1146/annurev.biochem.75.103004.142443

35. Pegg AE, Xiong H, Feith DJ, Shantz LM. 1998. S-adenosylmethionine decarboxylase: structure, function and regulation by polyamines. Biochem Soc Trans 26:580–586. https://doi.org/10.1042/bst0260580.

36. Yu TY, Schaefer J. 2008. REDOR NMR characterization of DNA packaging in bacteriophage T4. J Mol Biol 382:1031–1042. https://doi.org/10.1016/j.jmb.2008.07.077.

37. Crummett LT, Puxty RJ, Weihe C, Marston MF, Martiny JBH. 2016. The genomic content and context of auxiliary metabolic genes in marine cyanomyoviruses. Virology 499:219–229. https://doi.org/10.1016/j.virol.2016.09.016.

38. Clokie MR, Millard AD, Mann NH. 2010. T4 genes in the marine ecosystem: studies of the T4-like cyanophages and their role in marine ecology. Virol J 7:291. https://doi.org/10.1186/1743-422X-7-291.

39. Ignacio-Espinoza JC, Sullivan MB. 2012. Phylogenomics of T4 cyanophages: lateral gene transfer in the ‘core’ and origins of host genes. Environ Microbiol 14:2113–2126. https://doi.org/10.1111/j.1462-2920.2012.02704.x.

40. Chénard C, Chan AM, Vincent WF, Suttle CA. 2015. Polar freshwater cyanophage S-EIV1 represents a new widespread evolutionary lineage of phages. ISME J 9:2046–2058. https://doi.org/10.1038/ismej.2015.24.

41. Esposito D, Scocca JJ. 1997. The integrase family of tyrosine recombinases: evolution of a conserved active site domain. Nucleic Acids Res 25:3605–3614. https://doi.org/10.1093/nar/25.18.3605.

42. Nunes-Düby SE, Kwon HJ, Tirumalai RS, Ellenberger T, Landy A. 1998. Similarities and differences among 105 members of the Int family of site-specific recombinases. Nucleic Acids Res 26:391–406. https://doi.org/10.1093/nar/26.2.391.

43. Williams KP. 2002. Integration sites for genetic elements in prokaryotic tRNA and tmRNA genes: sublocation preference of integrase subfamilies. Nucleic Acids Res 30:866–875. https://doi.org/10.1093/nar/30.4.866.

44. Campbell A. 2003. Prophage insertion sites. Res Microbiol 154:277–282. https://doi.org/10.1016/S0923-2508(03)00071-8.

45. Paul JH. 2008. Prophages in marine bacteria: dangerous molecular time bombs or the key to survival in the seas? ISME J 2:579–589. https://doi.org/10.1038/ismej.2008.35.

46. Howard-Varona C, Hargreaves KR, Abedon ST, Sullivan MB. 2017. Lysogeny in nature: mechanisms, impact and ecology of temperate phages. ISME J 11:1511–1520. https://doi.org/10.1038/ismej.2017.16.

47. Bondy-Denomy J, Qian J, Westra ER, Buckling A, Guttman DS, Davidson AR, et al. 2016. Prophages mediate defense against phage infection through diverse mechanisms. ISME J 10:2854–2866. https://doi.org/10.1038/ismej.2016.79.

48. Casjens SR, Hendrix RW. 2015. Bacteriophage lambda: Early pioneer and still relevant. Virology 479–480:310–330. https://doi.org/10.1016/j.virol.2015.02.010.

49. Miller RV, Day M. 2008. Contribution of lysogeny, pseudolysogeny, and starvation to phage ecology. In: Abedon S (ed), Bacteriophage Ecology 114–143. Cambridge University Press.

50. Jiang SC, Paul JH. 1998. Significance of lysogeny in the marine environment: studies with isolates and a model of lysogenic phage production. Microb Ecol 35:235–243. https://doi.org/10.1007/s002489900079.

51. Maurice CF, Bouvier T, Comte J, Guillemette F, Giorgio PA. 2010. Seasonal variations of phage life strategies and bacterial physiological states in three northern temperate lakes. Environ Microbiol 12:628–641. https://doi.org/10.1111/j.1462-2920.2009.02103.x.

52. Williamson SJ, Houchin LA, McDaniel L, Paul JH. 2002. Seasonal variation in lysogeny as depicted by prophage induction in Tampa Bay, Florida. Appl Environ Microbiol 68:4307–4314. https://doi.org/10.1128/AEM.68.9.4307-4314.2002.

53. Payet J, Suttle CA. 2013. To kill or not to kill: The balance between lytic and lysogenic viral infection is driven by trophic status. Limnol Oceanogr 58:465–474. https://doi.org/10.4319/lo.2013.58.2.0465.

54. Wilson WH, Mann NH. 1997. Lysogenic and lytic production in marine microbial communities. Aquat Microb Ecol 13:95–100. https://doi.org/10.3354/ame013095.

55. Knowles B, Bailey B, Boling L, Breitbart M, Cobián-Güemes A, Del Campo J, et al. 2017. Variability and host density independence in inductions-based estimates of environmental lysogeny. Nat Microbiol 2:17064. https://doi.org/10.1038/nmicrobiol.2017.64.

56. Giovannoni SJ, Tripp HJ, Givan S, Podar M, Vergin KL, Baptista D, et al. 2005. Genome streamlining in a cosmopolitan oceanic bacterium. Science 309:1242–1245. https://doi.org/10.1126/science.1114057.

57. Grote J, Thrash JC, Huggett MJ, Landry ZC, Carini P, Giovannoni SJ, Rappé MS. 2012. Streamlining and core genome conservation among highly divergent members of the SAR11 clade. mBio 3:e00252–12. https://doi.org/10.1128/mBio.00252-12.

58. Thrash JC, Temperton B, Swan BK, Landry ZC, Woyke T, DeLong EF, Stepanauskas R, Giovannoni SJ. 2004. Single-cell enabled comparative genomics of a deep ocean SAR11 bathytype. ISME J 8:1440–1451. https://doi.org/10.1038/ismej.2013.243.

59. Giovannoni SJ, Thrash JC, Temperton B. 2014. Implications of streamlining theory for microbial ecology. ISME J 8:1553–1565. https://doi.org/10.1038/ismej.2014.60.

60. Touchon M, Bernheim A, Rocha EP. 2016. Genetic and life-history traits associated with the distribution of prophages in bacteria. ISME J 10:2744–2754. https://doi.org/10.1038/ismej.2016.47.

61. Shao Q, Trinh JT, Mcintosh CS, Christenson B, Balázsi G, Zeng L. 2017. Lysis- lysogeny coexistence: prophage integration during lytic development. Microbiologyopen 6:e00395. https://doi.org/10.1002/mbo3.395.

62. Feiner R, Argov T, Rabinovich L, Sigal N, Borovok I, Herskovits AA. 2015. A new perspective on lysogeny: prophages as active regulatory switches of bacteria. Nat Rev Microbiol 13:641–650. https://doi.org/10.1038/nrmicro3527.

63. Carini P, Steindler L, Beszteri S, Giovannoni SJ. 2013. Nutrient requirements for growth of the extreme oligotroph ‘Candidatus Pelagibacter ubique’ HTCC1062 on a defined medium. ISME J 7:592–602. https://doi.org/10.1038/ismej.2012.122.

64. Stingl U, Tripp HJ, Giovannoni SJ. 2007. Improvements of high-throughput culturing yielded novel SAR11 strains and other abundant marine bacteria from the Oregon coast and the Bermuda Atlantic Time Series study site. ISME J 1:361–37. https://doi.org/10.1038/ismej.2007.49.

65. Suttle CA, Fuhrman JA. 2010. Enumeration of virus particles in aquatic or sediment samples by epifluorescence microscopy. In Wilhelm S, Weinbauer M, Suttle C (eds) Manual of Aquatic Viral Ecology. American Society of Limnology and Oceanography: Waco, TX, USA pp 145–153. https://doi.org/10.4319/mave.2010.978-0-9845591-0-7.145.

66. Sambrook J, Russell DW. (eds). 2001. Molecular Cloning: A Laboratory Manual. Cold Spring Harbor, NY, USA: Cold Spring Harbor Laboratory.

67. Delcher AL, Harmon D, Kasif S, White O, Salzberg SL. 1999. Improved microbial gene identification with GLIMMER. Nucleic Acids Res 27:4636–4641. https://doi.org/10.1093/nar/27.23.4636

68. Aziz RK, Bartels D, Best AA, DeJongh M, Disz T, Edwards RA, et al. 2008. The RAST Server: Rapid Annotations using Subsystems Technology. BMC Genomics 9:75. https://doi.org/10.1186/1471-2164-9-75.

69. Lukashin AV, Borodovsky M. 1998. GeneMark.hmm: new solutions for gene finding. Nucleic Acids Res 26:1107–1115. https://doi.org/10.1093/nar/26.4.1107.

70. Lowe TM, Eddy SR. 1997. tRNAscan-SE: a program for improved detection of transfer RNA genes in genomic sequence. Nucleic Acids Res 25:955–964.

71. Edgar RC. 2004. MUSCLE: multiple sequence alignment with high accuracy and high throughput. Nucleic Acids Res 32:1792–1797. https://doi.org/10.1093/nar/gkh340.

72. Castresana J. 2000. Selection of conserved blocks from multiple alignments for their use in phylogenetic analysis. Mol Biol Evol 17:540–552. https://doi.org/10.1093/oxfordjournals.molbev.a026334.

73. Abascal F, Zardoya R, Posada D. 2005. ProtTest: selection of best-fit models of protein evolution. Bioinformatics 21:2104–2105. https://doi.org/10.1093/bioinformatics/bti263.

74. Stamatakis A. 2014. RAxML version 8: a tool for phylogenetic analysis and post-analysis of large phylogenies. Bioinformatics 30:1312–1313. https://doi.org/10.1093/bioinformatics/btu033.

75. Li L, Stoeckert CJ J, Roos DS. 2003. OrthoMCL: identification of ortholog groups for eukaryotic genomes. Genome Res 13:2178–2189. https://doi.org/10.1101/gr.1224503.

76. van Dongen S, Abreugoodger C. 2012. Using MCL to extract clusters from networks. Methods Mol Biol 804:281–295. https://doi.org/10.1007/978-1-61779-361-5_15.

